# Gut Bacteria Metabolize Natural and Synthetic Steroid Hormones via the Reductive OsrABC Pathway

**DOI:** 10.1101/2024.10.08.617280

**Authors:** Christian Jacoby, Kaylie Scorza, Lia Ecker, Paola Nol Bernardino, Alexander S. Little, Mary McMillin, Ramanujam Ramaswamy, Anitha Sundararajan, Ashley M. Sidebottom, Huaiying Lin, Keith Dufault-Thompson, Brantley Hall, Xiaofang Jiang, Samuel H. Light

**Affiliations:** Duchossois Family Institute, University of Chicago, Chicago, IL 60637, USA; Department of Microbiology, University of Chicago, Chicago, IL 60637, USA; National Library of Medicine, National Institutes of Health, Bethesda, MD 20894, USA; Department of Cell Biology and Molecular Genetics, University of Maryland, College Park, College Park, MD 20742, USA

## Abstract

Steroid hormone metabolism by the gut microbiome affects host physiology, but the underlying microbial pathways remain incompletely understood. Here, we isolate a novel human gut bacterium, *Clostridium steroidoreducens*^T^ strain HCS.1 that reduces cortisol and related steroid hormones to 3β,5β-tetrahydrosteroid products. Through transcriptomics and enzymatic discovery, we establish the *C. steroidoreducens* OsrABC steroid hormone pathway. OsrA is a 3-oxo-Δ^1^-steroid hormone reductase that targets synthetic glucocorticoids, including prednisolone— a frontline Crohn’s disease therapy. OsrB is a 3-oxo-Δ^4^-steroid reductase that converts steroid hormones to 5β-dihydrosteroid intermediates, which OsrC subsequently reduces to 3β,5β-tetrahydro products. Homologs of *osrA* and *osrB* predict steroid-reducing activity across gut bacteria and are enriched in Crohn’s disease patient metagenomes. Consistent with a role in modulating drug efficacy, *C. steroidoreducens* colonization decreases prednisolone bioavailability in gnotobiotic mice. These findings thus define a previously unrecognized pathway for microbial steroid hormone inactivation and establish a mechanistic basis for bacterial interference with anti-inflammatory therapies.

## Introduction

Steroid hormones encompass a broad class of biologically active molecules that play crucial roles in diverse physiological processes. Glucocorticoids, like cortisol, are involved in regulating inflammation, immune response, and metabolism,^1^ while sex steroids, including estrogens, androgens, and progestins, regulate reproductive functions and secondary sexual characteristics.^2^ Due to their wide-ranging effects, both natural and synthetic steroid hormones are commonly used in medical therapies to manage conditions such as autoimmune diseases, hormone deficiencies, and cancers.

The gut microbiome mediates diverse phenotypes through the modification of host- and diet-derived metabolites, including various steroids. Research on microbiome steroid metabolism has primarily focused on bile acids, products of which are important for multiple host phenotypes.^3-5^ However, while less studied, steroid hormones have also been identified as an important class of substrates in the gut. These molecules interact with gut microbes after entering the gastrointestinal tract via therapeutic oral or rectal administration or through excretion in bile.^6^ Bile is likely a particularly important source of intestinal steroid hormones, as 9-22% of endogenous cortisol,^7,8^ 10-15% of testosterone,^9^ 20-30% of corticosterone,^10^ and up to 30% of progesterone^11^ are eliminated from the body through this route.

Gut microbes generate multiple products from steroid hormones, including cortisol and corticosterone derivatives that serve as fecal biomarkers for stress in animal research studies.^12,13^ In addition to passing into feces, intestinal microbial steroid hormone products can reenter the bloodstream through enterohepatic circulation. In humans, this is evidenced by the rectal administration of cortisol leading to an increase in specific circulating cortisol derivatives in a microbiome-dependent manner.^14,15^ Other studies provide evidence that microbial products of progesterone metabolism similarly enter systemic circulation.^6,16^

The impact of microbial steroid hormone metabolism has been linked to several aspects of mammalian biology. Microbial inactivation of orally administered steroids, including side-chain cleavage of synthetic glucocorticoids, reduces the bioavailability of these drugs.^17,18^ Microbial pathways that dehydroxylate corticosterone and convert steroid precursors to androgens generate metabolites that contribute to hypertension in animal models^19,20^ and promote castration-resistant prostate cancer.^21-23^ Bacterial metabolisms that alter the concentration of steroid hormones with distinct biological activities thus have diverse consequences for mammalian biology.

Previous studies have reported that some gut bacteria reduce the Δ^4^-bond in steroid hormones, generating 5β-steroid derivatives.^24^ This activity decreases the anti-inflammatory and androgenic properties of glucocorticoids and androgens, respectively, and converts progestins into a neuroactive form.^25^ While it stands to reason that Δ^4^-steroid hormone reduction may have implications for host biology, the molecular basis and the broader significance of activities in the gut microbiome remains unknown.

Here, we describe the isolation and characterization of a novel steroid hormone-metabolizing gut bacterium, *Clostridium steroidoreducens* HCS.1. By employing a multidisciplinary approach, integrating genomics, transcriptomics, and metabolomics, we characterize the *osrABC* reductive steroid hormone pathway. These findings provide new insights into the diversity of steroid hormone metabolism in the gut microbiome and its potential impact on host health.

## Results

### *Clostridium steroidoreducens* is a novel steroid hormone-enriched gut bacterium

To select for members of the gut microbiome with metabolic capabilities that provide a selective advantage in the presence of steroid hormones, we passaged a fecal sample from a healthy human donor with no history of inflammatory disease in a nutritionally limited base medium supplemented with individual glucocorticoids (cortisol or corticosterone) or sex steroids (progesterone or testosterone) (**Figure 1A**). We isolated strains from the final enrichment passages and cultivated them on steroid hormone-infused solid media. We identified one strain, HCS.1, that cleared insoluble cortisol or progesterone from the solid media and accumulated a white precipitate indicative of a potential reaction product at colony centers on progesterone-infused media (**Figure 1B and 1C**). Consistent with HCS.1 possessing a selective advantage in the presence of steroid hormones, 16S rRNA amplicon sequencing of the final enrichment passages revealed that an amplicon sequence variant matching the HCS.1 16S rRNA sequence was enriched from below the limit of detection in the fecal inoculum to 0.7-23.6% of the microbial community following cortisol, corticosterone, progesterone, or testosterone enrichment (**Figure 1D**).

**Figure 1.**
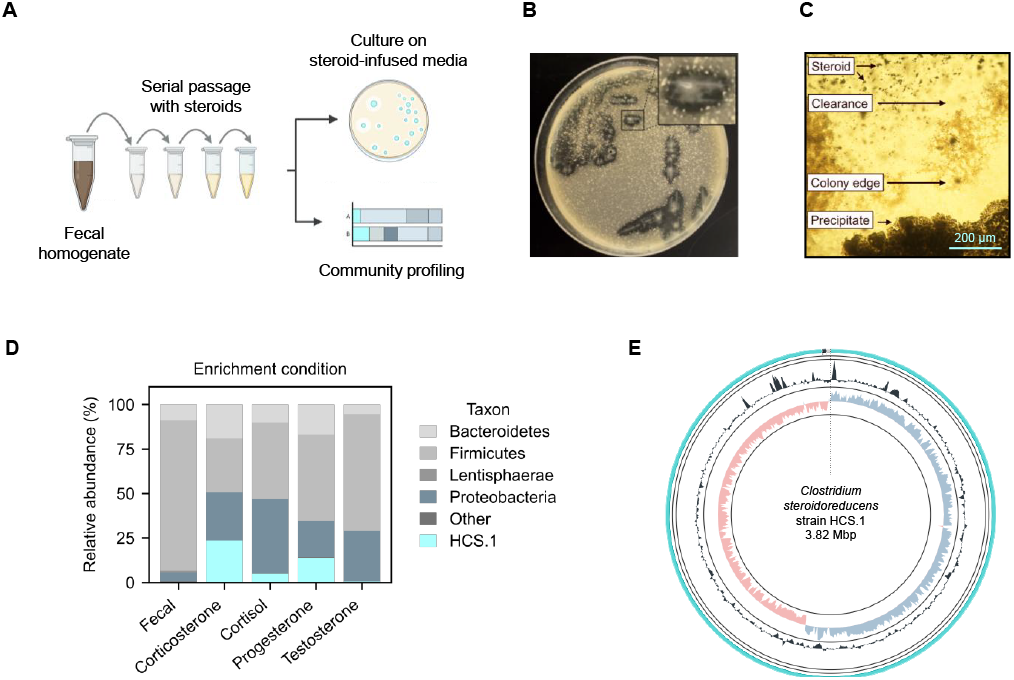
*Clostridium steroidoreducens* HCS.1 is a novel steroid-enriched species. (A) Schematic overview of steroid enrichment and strain isolation experiments. (B) HCS.1 strain colonies on cortisol-infused media. (C) HCS.1 strain colonies on progesterone-infused media. (D) Microbial community profile of the final steroid hormone enrichment passage based on 16S rRNA amplicon sequencing. (E) Circular representation of the HCS.1 genome.

To facilitate further strain characterization, we sequenced and assembled HCS.1 DNA into 3 circularized contigs, comprising a genome (NCBI accession: CP170704) and two plasmids (NCBI accession: CP170705, CP170706), that contain 3,816,008 base pairs with 28.4% G + C content, 3595 protein-coding genes, and 111 RNA genes (**Figure 1E**). Phylogenetic analyses revealed that HCS.1 was closely related to *Clostridium chrysemydis* but represents a novel species of the genus *Clostridium*, based on an average nucleotide identity (ANI) of 92.09% to the closest reference genome and accepted taxonomic assignment criteria (**Figure S1**).^26^ In recognition of its steroid hormone reductase activities detailed below, we assigned HCS.1 the species name *Clostridium steroidoreducens*.

### *C. steroidoreducens* possesses broad steroid hormone reductase activity

To determine whether the *steroidoreducens* HCS.1 steroid clearance phenotype was indicative of metabolic activity, we employed an LC-MS-based assay to track the fate of cortisol in *C. steroidoreducens* HCS.1 culture. We observed that cortisol was fully depleted from the media, coinciding with the emergence of a minor and a major product. By comparison with compound reference standards, we inferred that the major and minor products corresponded to 5β-dihydro- and 3β,5β-tetrahydrocortisol, respectively (**Figure 2A**).

**Figure 2.**
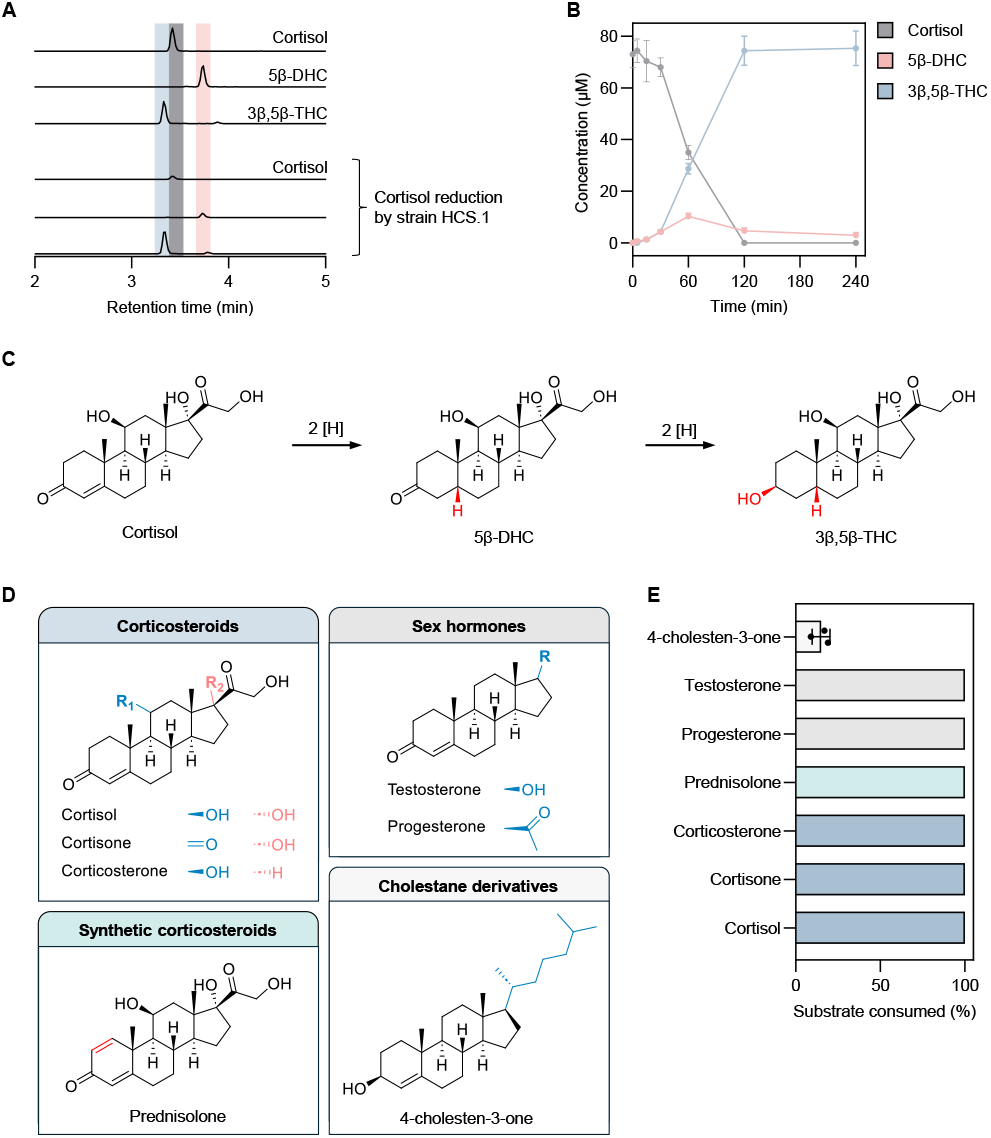
*C. steroidoreducens* HCS.1 possesses promiscuous 3-oxo-Δ^4^-beta steroid hormone reductase activity. (A) Products formed from *C. steroidoreducens* HCS.1 incubation with cortisol. (B) Time-course analysis of cortisol metabolism by *C. steroidoreducens* HCS.1. Error bars refer to the standard deviation of three biological replicates (C) Proposed pathway for cortisol reduction by *C. steroidoreducens* HCS.1. DHC and THC stand for dihydrocortisol and tetrahydrocortisol, respectively. (D) Steroid substrates tested for *C. steroidoreducens* HCS.1. (E) Measured *C. steroidoreducens* HCS.1 steroid substrate consumption. Error bars refer to the standard deviation of three independent replicates.

Tracking *C. steroidoreducens* HCS.1 cortisol metabolism over time, we observed that 5β-dihydrocortisol transiently accumulated, peaking at 60 minutes before decreasing to less than 2% of the total glucocorticoid present by 120 minutes (**Figure 2B**). By contrast, 3β,5β-tetrahydrocortisol steadily accumulated following the introduction of cortisol, exceeding 98% of the glucocorticoid present by 120 minutes (**Figure 2B**). These results suggest that, in contrast to previously characterized bacterial steroid dehydroxylation^27^ and side chain-cleaving^28^ activities, *C. steroidoreducens* HCS.1 exclusively reduces cortisol, converting it to 3β,5β-tetrahydrocortisol via a 5β-dihydrosteroid intermediate (**Figure 2C**).

To address the specificity of *C. steroidoreducens* HCS.1 steroid utilization, we next tested the strain’s activity on a panel of steroids with variable functional groups at multiple positions on the sterol core (**Figure 2D**). We found that *C. steroidoreducens* tolerated substitutions at C1, C11, and C17 positions, exhibiting activity on distinct glucocorticoids (corticosterone, cortisone, prednisolone) and sex steroids (progesterone, testosterone) (**Figure 2E**). In contrast to these more polar steroids, the hydrophobic cholesterol derivative 4-cholesten-3-one was a poor substrate (**Figure 2E**). These results establish *C. steroidoreducens* as a steroid hormone-reducing gut bacterium with broad substrate specificity.

### OsrB is a Fe-S flavoenzyme family 3-oxo-Δ^4^-steroid hormone reductase

We next sought to identify the mechanism of steroid hormone reduction by *C. steroidoreducens*. As bacterial reductases are often induced in the presence of their substrate,^27,29^ we employed a transcriptomics-based approach to identify candidate steroid hormone reductases in *C. steroidoreducens*. We performed RNA-seq analysis on *C. steroidoreducens* cells cultivated in the presence or absence of cortisol and identified 30 genes that were induced >2-fold when cortisol was present (**Dataset S1**). Two of the most highly induced genes, which we renamed *osrA* (NCBI accession MFU7516964.1) and *osrB* (NCBI accession MFU7517346.1) for oxosteroid reductase A and B, encoded proteins annotated as *fadH*-like 2,4-dienoyl-CoA reductases (**Figure 3A, Dataset S1**).

**Figure 3.**
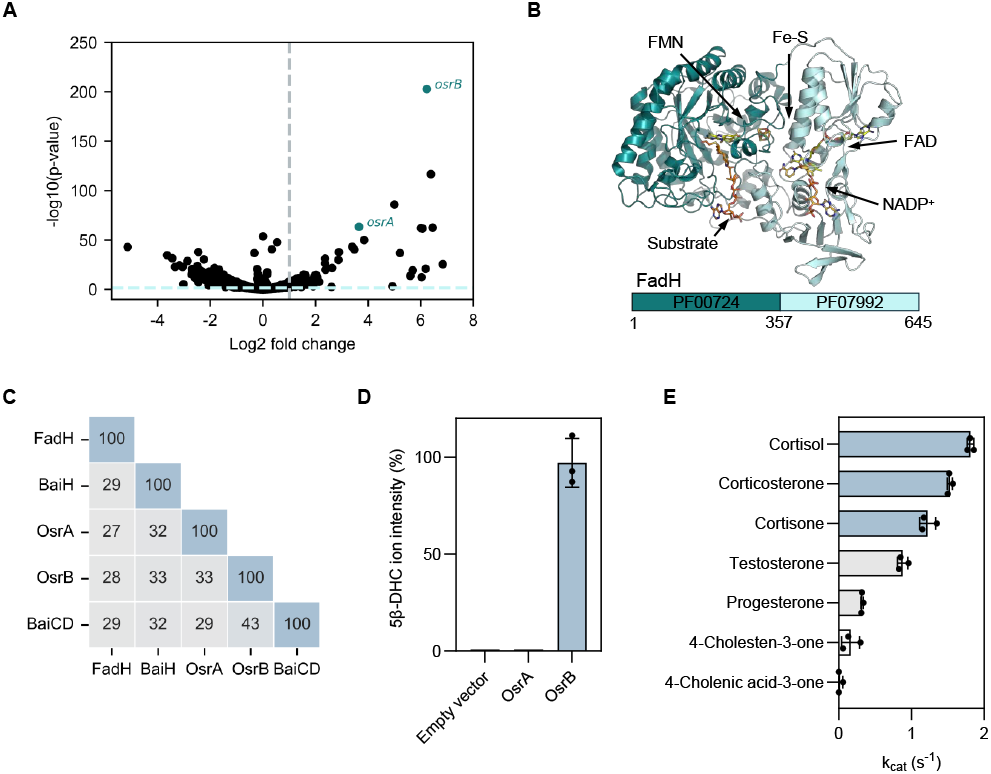
OsrB is a 3-oxo-Δ^4^-steroid hormone reductase. (A) Gene expression of *C. steroidoreducens* HCS.1 in the presence versus absence of cortisol. Gray and red dashed lines indicate genes with statistical significance and >2-fold induction in response to cortisol, respectively. (B) Crystal structure of Fe-S flavoenzyme 2,4-dienoyl-CoA reductase (FadH) bound to ligands (PDB code: 1PS9). (C) Percent sequence identity of OsrA and OsrB to Fe-S flavoenzymes FadH and bile acid reductases BaiH and BaiCD. (D) Conversion of cortisol to 5β-dihydrocortisol (5β-DHC) by *E. coli* expressing *osrA* or *osrB* versus an empty vector control. Error bars refer to the standard deviation of three independent replicates. (E) Rate of reduction of indicated steroid hormones by purified OsrB. Error bars refer to the standard deviation of three independent replicates.

*E. coli* 2,4-dienoyl-CoA reductase is the best characterized member of the “Fe-S flavoenzyme family” of oxidoreductases that contain a conserved N-terminal substrate-binding domain (PF00724) and a C-terminal NAD(P)H cofactor-binding domain (PF07992) (**Figure 3B**).^30^ Divergent members of the Fe-S flavoenzyme family are widespread in gut bacteria and possess distinct substrate specificities for host- and diet-derived metabolites.^31^ Consistent with OsrA and OsrB representing novel Fe-S flavoenzyme subtypes with distinct substrates, we observed that these enzymes exhibited remote sequence homology to previously characterized Fe-S flavoenzymes, including *Clostridium scindens* Fe-S flavoenzymes, BaiCD and BaiH, which reduce bile acid intermediates structurally related to steroid hormones (**Figure 3C**).^30,32^

To test steroid reductase activity of OsrA and OsrB, we heterologously produced the enzymes in anaerobically cultured *E. coli* cells. We found cells expressing *osrB*, but not *osrA*, reduced cortisol to 5β-dihydrocortisol (**Figure 3D**). Studies with anaerobically purified OsrB revealed that NADH and NADPH were poor electron donors for OsrB, suggesting the enzyme requires alternative reducing equivalents. Using the artificial electron donor methyl viologen, we observed that OsrB efficiently reduced a variety of steroid hormones, while exhibiting only minor activity toward cholesterol and no detectable activity with bile acid structural analogs (**Figure 3E**). These results thus establish OsrB as a promiscuous 3-oxo-Δ^4^-steroid hormone reductase that uses a presently unidentified electron donor.

### OsrC is a short chain dehydrogenase family 3-oxo-5β-steroid hormone oxidoreductase

We next sought to identify the *C. steroidoreducens* enzyme responsible for reduction of the 3-oxo group on the 5β-dihydrosteroid intermediate generated by OsrB. Microbial bile acid oxidoreductases with specificity for 3α-, 3β-, 7α-, 7β-, 12α- and 12β-hydroxyl groups have been previously identified.^33-36^ As these characterized steroid oxidoreductases are members of the short chain dehydrogenase (SDR) enzyme superfamily, we reasoned the *C. steroidoreducens* enzyme was likely related to this family. An analysis of the *C. steroidoreducens* genome identified 12 genes with SDR domains. However, none were induced by cortisol or exhibited high sequence similarity to previously characterized bile acid oxidoreductases, suggesting that the gene encoding this enzyme is constitutively expressed.

As these analyses failed to identify obvious candidates, we adapted a previously described approach to perform an unbiased screen of SDR-containing *C. steroidoreducens* proteins for 3-oxo-5β-steroid hormone reductase activity.^37^ We confirmed soluble expression of all 12 SDRs in *E. coli* and tested the activity of overexpressing *E. coli* strains on 5β-dihydrocortisol (**Figure 4A, Figure S2**). We identified two SDRs (ACFYH6_12520 and ACFYH6_05340) that produced 3β,5β-tetrahydrocortisol and two others (ACFYH6_12160 and ACFYH6_06930) that yielded 3α,5β-tetrahydrocortisol (**Figure 4A**).

**Figure 4.**
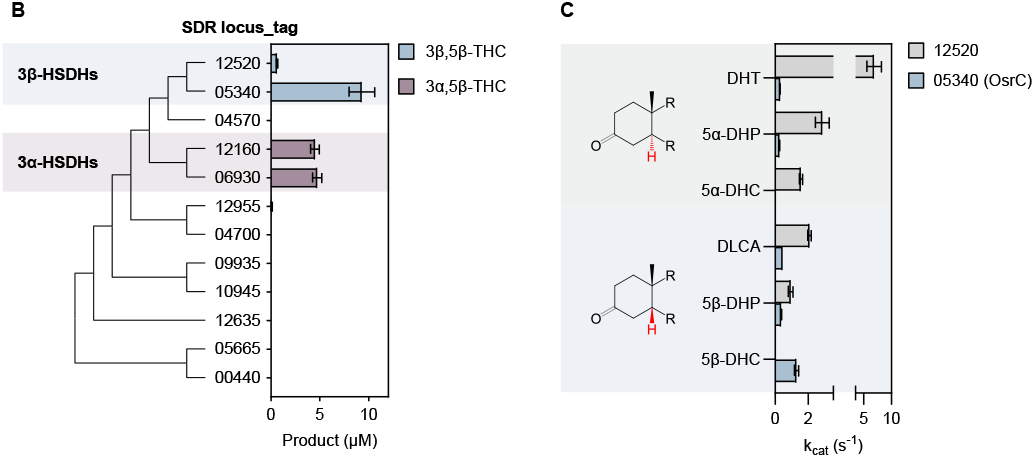
OsrC is a 3-oxo-5β-steroid hormone oxidoreductase. (A) Phylogenetic analysis of twelve C. *steroidoreducens* HCS.1 SDR superfamily enzymes, labeled by numeric identifiers (XXXXX) corresponding to their locus tag suffixes (ACFYH6_XXXXX). Product formed from 5β-dihydrocortisol by *E. coli* strains overexpressing SDR domain-containing proteins are shown with their respective gene identifiers. Error bars refer to the standard deviation of three independent replicates. (B) Kinetic parameters of reduction of indicated steroid hormones by purified SDR domain-containing proteins. Abbreviations stand for tetrahydrocortisol (THC), dihydrotestosterone (DHT), dihydroprogesterone (DHP), dihydrocortisol (DHC), dehydrolithocholic acid (DLCA) and hydroxysteroid dehydrogenase (HSDH).

Studies with aerobically purified ACFYH6_12520 and ACFYH6_05340 revealed both enzymes preferred NADPH as an electron donor but possessed distinct specificities for steroid substrates. ACFYH6_12520 showed a pronounced preference for 5α-steroids and exhibited weak activity with 5β-steroid substrates (**Figure 4B**). Conversely, ACFYH6_05340 displayed a preference for 5β-steroid hormones and tolerated various functional groups at the C9 or C17 positions (**Figure 4B**). We further found that ACFYH6_05340 exhibited a preference for steroid hormones relative to the comparable bile acid derivative lithocholic acid, with the highest activity observed towards 5β-dihydrocortisol. We thus conclude that ACFYH6_05340 is a 3-oxo-Δ^4^-steroid hormone reductase and, on this basis, renamed it OsrC (NCBI accession MFU7515415.1).

### OsrA is a Fe-S flavoenzyme family 3-oxo-Δ^1^-steroid hormone reductase active on synthetic glucocorticoids

Synthetic glucocorticoids possess potent anti-inflammatory properties and are widely used to treat a range of pathologies, including inflammatory bowel disease (IBD).^38^ The synthetic glucocorticoids dexamethasone, prednisone, prednisolone, budesonide and methylprednisolone contain a Δ^1^-double bond that is absent in natural glucocorticoids and which significantly extends their half-life (**Figure 5A**).^39^ As our initial screen of steroids identified prednisolone as a substrate for *C. steroidoreducens* (**Figure 2E**), we sought to address the molecular basis of synthetic glucocorticoid metabolism. We first tested the activity of *C. steroidoreducens* on additional synthetic glucocorticoids (dexamethasone, prednisone, budesonide and methylprednisolone) and found that all were reduced to 3β,5β-tetrahydrocortisol derivatives, indicating that the bacterium possesses both Δ^1^- and Δ^4^-steroid hormone reductase activities (**Figure 5B**).

**Figure 5.**
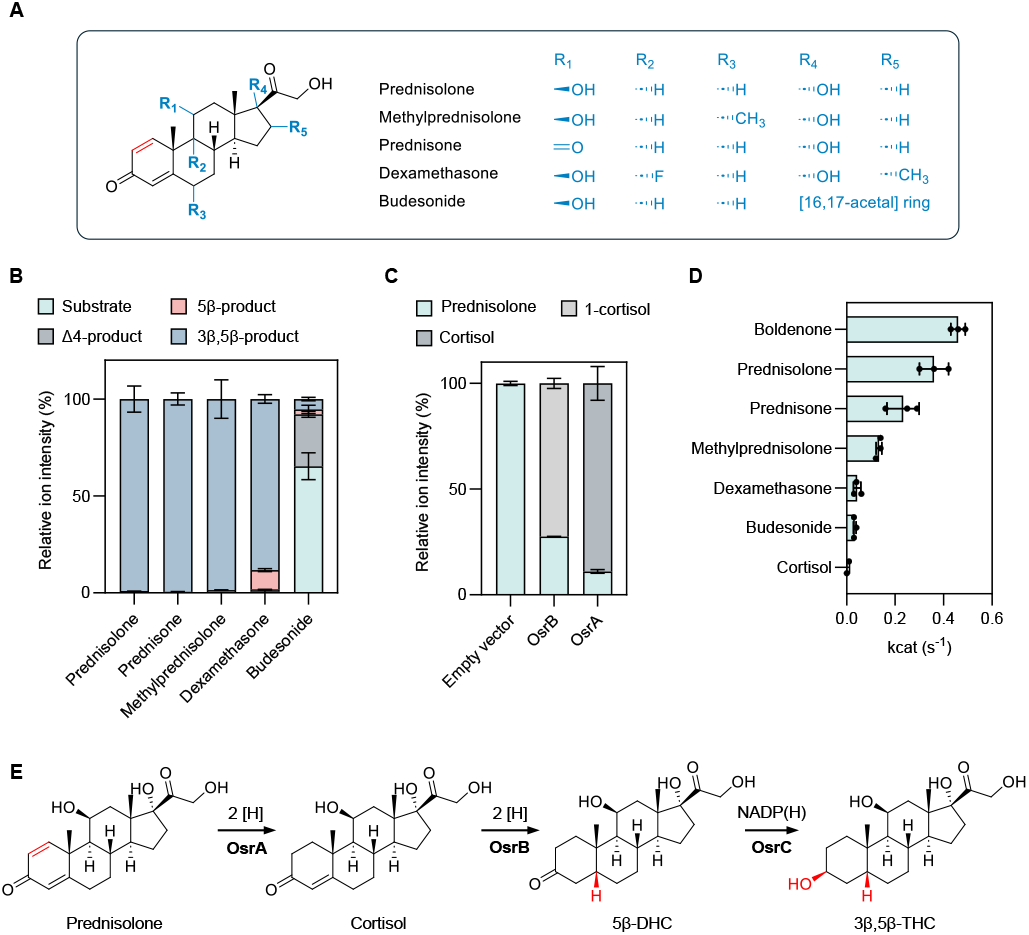
OsrA is a 3-oxo-Δ^1^-reductase essential for complete reduction of synthetic glucocorticoids. (A) Structure of synthetic glucocorticoids used in assays. (B) Products formed from synthetic glucocorticoids following incubation with *C. steroidoreducens* HCS.1 cells. Error bars refer to the standard deviation of three independent replicates. (C) Percent prednisolone conversion to cortisol following incubation of *E. coli* cells with *osrA*- and *osrB*-expressing plasmids or an empty vector control. Error bars refer to the standard deviation of three independent replicates. (D) Rate of reduction of indicated synthetic steroid hormones by purified OsrA. Error bars refer to the standard deviation of three independent replicates. (E) *C. steroidoreducens* HCS.1 steroid reduction pathway identified in this study.

Considering the similarity of the Δ^1^-reduction to the OsrB catalyzed Δ^4^-reduction, we next tested the activity of OsrA and OsrB and found that the two enzymes generated distinct cortisol and testosterone isomers from prednisolone and the synthetic androgen boldenone, respectively (**Figure 5C, Figure S3**). Based on comparison to reference standards, we establish that OsrA and OsrB products reflected Δ^1^- and Δ^4^-steroid hormone reductase activities, respectively (**Figure S3**). To validate the substrate specificity of OsrA, we anaerobically purified the enzyme and conducted biochemical assays with defined electron donors. Kinetic analyses revealed that while both methyl viologen and NADH supported enzymatic activity, methyl viologen yielded markedly higher reaction rates. Substrate profiling demonstrated that purified OsrA efficiently reduced all tested synthetic glucocorticoids but lacked detectable activity toward cortisol, clearly demonstrating the specificity of OsrA for the Δ^1^ double bond present exclusively in synthetic steroids (**Figure 5D**). These results demonstrate that OsrA functions as a Δ^1^-steroid hormone reductase that acts in conjunction with OsrB and OsrC to reduce synthetic steroid hormones to 3β,5β-reduced products (**Figure 5E**).

### Steroid hormone reductase activities are common in gut bacteria and correlate with the distribution of *osrA* and *osrB* homologs

We next sought to address the breadth of steroid hormone reductase activity in the gut microbiome. We performed BLASTp searches of OsrA, OsrB, and OsrC in the Unified Human Gastrointestinal Genome catalog of representative genomes and metagenome-assembled genomes, which includes 4,644 prokaryotic species that colonize the human gastrointestinal tract.^40^ These searches identified homologs with high sequence homology to OsrA, OsrB, and OsrC in 2, 59, and 90 genomes, respectively (**Dataset S2**). Genomes encoding *osrABC* homologs included gram-positive bacterial species from multiple taxa, primarily from the Erysipelotrichaceae and Lachnospiraceae families.

To determine the association of *osrABC* homologs with observed *C. steroidoreducens* phenotypes, we selected 117 bacterial strains capturing the taxonomic diversity of the human gut microbiome for experimental characterization. We assayed these strains on solid media for steroid clearance/precipitate accumulation and further tested a subset for cortisol and prednisolone activity. From these studies, we identified 29 strains from 14 species with a steroid clearance/precipitate accumulation phenotype and isolates from 6 species exhibiting steroid hormone reductase activity (**Figure 6A-6C and Dataset S3**).

**Figure 6.**
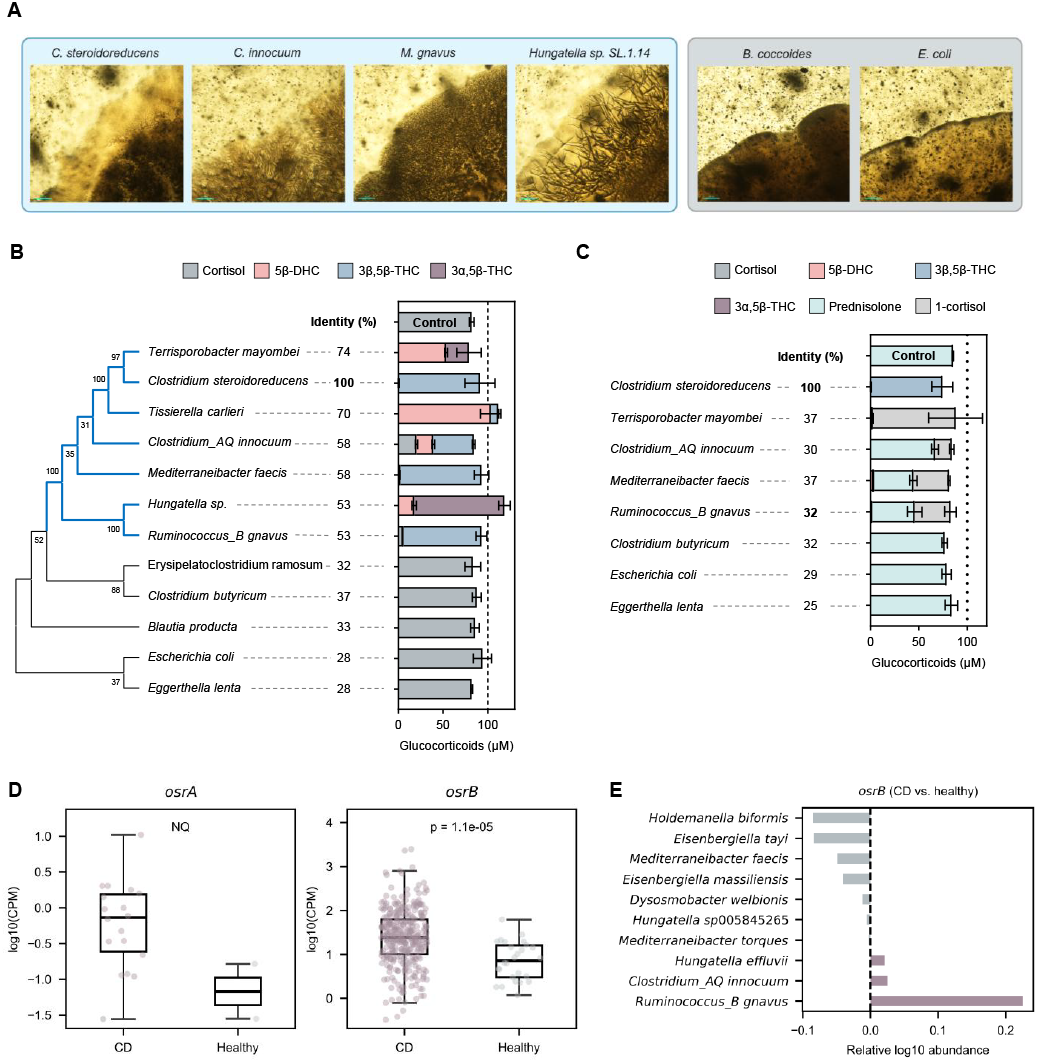
Steroid reductase activities are widespread in gut bacteria and elevated in active Crohn’s disease. (A) Representative images of gut bacteria grown on progesterone-infused media, showing steroid clearance/precipitate accumulation-positive (blue background) and -negative (gray background) colonies. Scale bar, 200 µm. (B) Glucocorticoids produced by gut bacterial isolates after incubation with cortisol. The protein with the highest sequence identity to OsrB encoded by each genome was used to generate the tree. Error bars refer to the standard deviation of three independent replicates. (C) Glucocorticoids produced by gut bacterial isolates after incubation with prednisolone. Identity refers to the sequence identity of the protein with the highest sequence identity to OsrA encoded by each genome. Error bars refer to the standard deviation of three independent replicates. (D) Read counts mapping to *osrA* and *osrB* homologs in metagenomes from healthy and Crohn’s disease (CD) patients. Only metagenomes with at least one read mapping to a gene are included in the analysis. Sample sizes: for osrA, *n*_Healthy_ = 2 and *n*_CD_ = 28; for osrB, *n*_Healthy_ = 25 and *n*_CD_ = 288. NQ refers to the not quantified statistical difference, due to the low number of healthy metagenomes with reads mapping to *osrA*. CPM refers to copies per million. (E) Difference in *osrB* homolog abundance for taxa showing the greatest changes in relative abundance between healthy and CD metagenomes.

Comparing strain genotypes to observed phenotypes revealed several patterns. First, the presence of an *osrB* homolog in a genome strongly predicted both steroid clearance/precipitate accumulation and steroid hormone Δ^4^-reductase activity (**Figure 6B and Dataset S3**). Second, the absence of *osrA* homologs tracked with a consistent lack of steroid hormone Δ^1^-reductase activity (**Figure 6C and Dataset S3**). Third, while the presence of an *osrC* homolog was associated with production of 3β,5β-tetrahydrocortisol, the absence of an *osrC* homolog was not predictive of the fate of the C3 functional group. Indeed, *osrC*-negative strains varied in their major cortisol product, generating either 5β-dihydrocortisol, 3α,5β-tetrahydrocortisol, or 3β,5β-tetrahydrocortisol (**Figure 6B and Dataset S3**). These results provide evidence that *osrA* and *osrB* specifically confer Δ^1^- and Δ^4^ -steroid hormone reductase activities, respectively, while *osrC* likely represents one of multiple evolutionarily distinct 3-oxo-5β-steroid hormone oxidoreductases.

### *osrB* is prevalent in human fecal metagenomes and associated with active Crohn’s disease

Having established the relevance of *osrABC* homologs for steroid reductase activity in gut bacteria, we next sought to determine the prevalence of the pathway in the human gut. We focused our analysis on *osrA* and *osrB*, since these homologs reliably predicted steroid hormone reductase activities of assayed strains. We recruited reads to *osrA* and *osrB* homologs in a collection of 1,491 previously published healthy human fecal metagenomes. We detected at least one read mapping to *osrA* and *osrB* homologs in 2.2% and >98.9% of samples, respectively (**Dataset S4**). Within most metagenomes, multiple *osrB* homologs recruited substantial read counts. By contrast, the majority of *osrA* reads mapped to *Clostridium tertium osrA* homologs, often with only one or two reads per metagenome (**Dataset S4**). These analyses demonstrate that *osrB* homologs are common in the gut but that *osrA* homologs are confined to bacteria that colonize the gut at a low relative abundance.

In mouse models of intestinal inflammation, local glucocorticoid production signals through myeloid and intestinal epithelial cells to suppress inflammation.^41-45^ A similar mechanism is likely important in humans, as both natural and synthetic glucocorticoid variants, including cortisol and prednisolone, are frontline treatments for IBD.^46,47^ Considering the significance of both endogenous production and therapeutic application of glucocorticoids in IBD, we reasoned that OsrA and OsrB activity could be clinically relevant in this patient population.

We analyzed 314 metagenomes from the Lewis et al.^48^ study of active Crohn’s disease patients, including a subset treated with glucocorticoids (**Dataset S5**). We observed *osrB* homologs were elevated in Crohn’s disease patient relative to a healthy control population (**Figure 6D**). Further scrutiny revealed that the increased abundance of *osrB* homologs from *Ruminococcus_B osrB gnavus*, homologs from *Clostridium_AQ innocuum*, two taxa previously associated with Crohn’s disease inflammation,^49,50^ was the primary driver of this association (**Figure 6E**).

We further found *osrA* homologs similarly exhibited elevated abundance in Crohn’s disease metagenomes, but their low prevalence coupled with the relatively small sample size of this dataset complicated statistical analysis of the significance of this relationship (**Figure 6D**). To address this issue, we expanded our dataset to include 1,537 metagenomes from multiple separate studies that included Crohn’s disease and control populations. Analysis of this larger dataset confirmed that *osrA* homologs were significantly elevated in Crohn’s disease patient metagenomes and revealed that the *osrA* from *Clostridium tertium* was the primary driver of this association (**Figure S4A and S4B, Dataset S5**).

To determine whether identified associations extended an independent dataset, we evaluated 569 Crohn’s disease patient metagenomes collected as part of the Integrative Human Microbiome Project (**Dataset S6**). We observed a similar association between elevated abundance of *osrA* and *osrB* homologs and microbiome dysbiosis scores used as a proxy for active Crohn’s disease in this study (**Figure S5A-S5C, Dataset S6**).^51^ Underscoring the relevance of these observations to active Crohn’s disease, metagenomes from this study with microbiome dysbiosis scores consistent with inactive Crohn’s disease exhibited intermediate *osrB* homolog levels between dysbiotic Crohn’s disease and control non-IBD populations (**Figure S5A, Dataset S6**).

We further investigated the relationship between microbial steroid reductases and glucocorticoid treatment in Crohn’s disease patients, as reported in the Lewis et al.^48^ study. Our analysis revealed that glucocorticoid therapy was associated with an increase in *osrA* homolog prevalence, as these homologs were detected in 9.6% of glucocorticoid-treated patients compared to only 2.4% in untreated patients (**Dataset S5**). Interestingly, in metagenomes containing *osrA* homologs, their abundance did not significantly differ between treated and untreated patients, suggesting that synthetic glucocorticoids may promote the colonization but not proliferation of bacteria capable of metabolizing these drugs (**Figure S6)**.

### Microbial steroid metabolism reduces prednisolone bioavailability in gnotobiotic mice

Considering that synthetic glucocorticoids represent frontline IBD therapies and that *osrA* and *osrB* homologs are enriched in Crohn’s disease metagenomes, we sought to clarify the effect of *C. steroidoreducens* colonization on prednisolone bioavailability and therapeutic exposure. We first validated that our analytical methods could resolve orally administered prednisolone and endogenous cortisol but not bacterial metabolites (dihydro- and tetrahydrocortisol products) in germ-free control mice (**Figure 7A**). To facilitate direct assessment of bacterial effects on glucocorticoid metabolism without confounding influences from complex microbial communities, we next colonized germ-free mice with *C. steroidoreducens* HCS.1 and administered prednisolone via oral gavage to mimic therapeutic dosing (**Figure 7B**).

**Figure 7.**
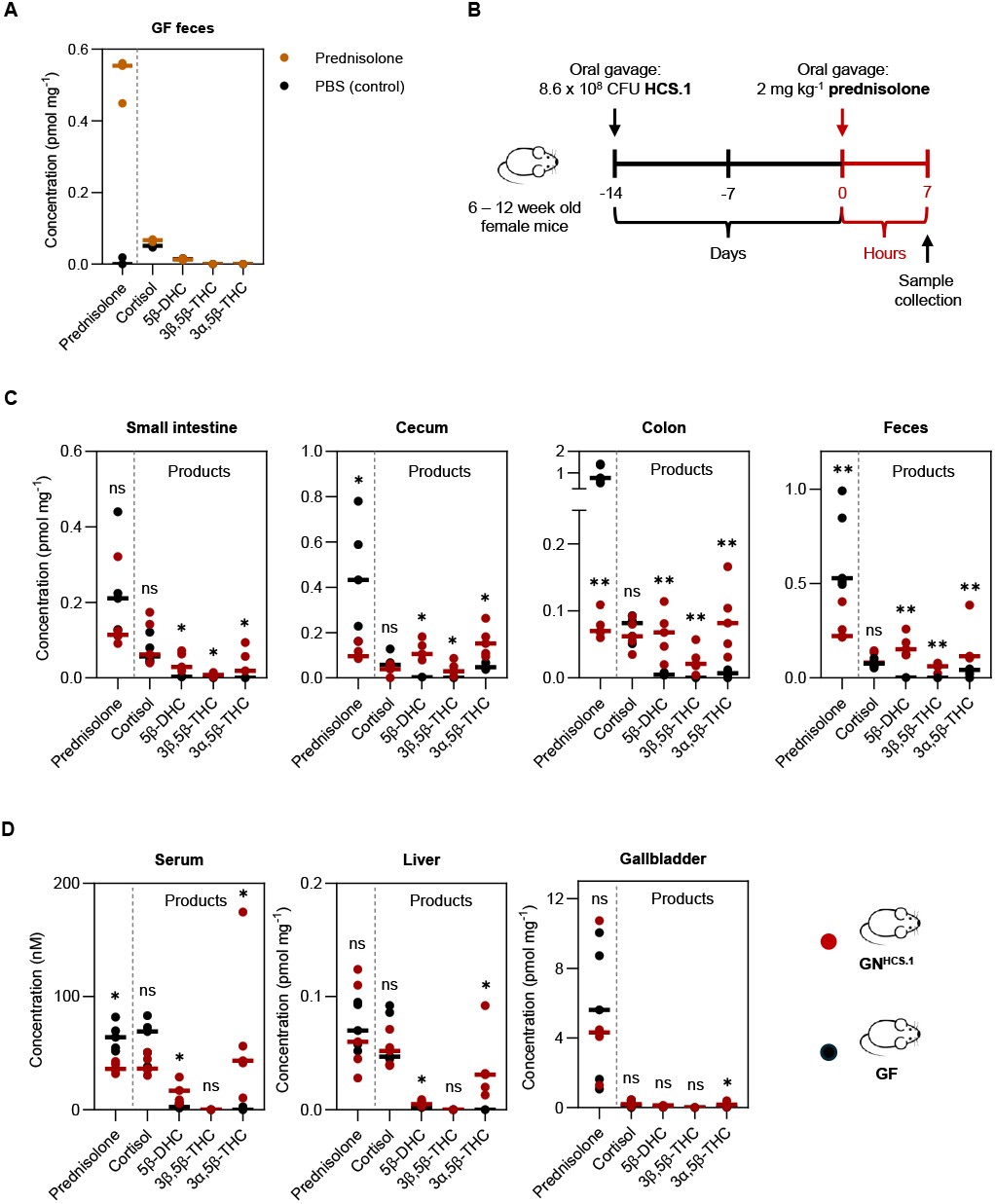
Mono-colonization with *Clostridium steroidoreducens* HCS.1 promotes glucocorticoid inactivation in gnotobiotic mice. (A) Glucocorticoid levels in feces of germfree mice treated with PBS (control) or 2 mg/kg prednisolone. (B) Timeline of gnotobiotic mouse colonization, glucocorticoid administration, and sampling. Colonization with *C. steroidoreducens* HCS.1 in mice was confirmed after 7 days by direct fecal culturing and PCR amplification of *osrB* from the resulting isolates. (C) Glucocorticoid levels detected in the small intestine, cecum, colon, and feces of germfree mice (n = 5) and *C. steroidoreducens*-mono-colonized mice (n = 5) seven hours after oral gavage with 2 mg/kg of prednisolone. (D) Glucocorticoid levels detected in serum, liver, and gallbladder. P-values were calculated using two-sided Mann–Whitney U tests, followed by false discovery rate (FDR) correction using the Benjamini–Hochberg method. Significance: p < 0.05 (\**), p < 0*.*01 (**), p < 0*.*001 (\*\*\**).

Consistent with *C. steroidoreducens* primarily acting in the lower gastrointestinal tract, LC-MS analysis of gastrointestinal contents revealed that prednisolone levels were unchanged in the small intestine but decreased 2-10-fold in the cecum, colon, and feces of *C. steroidoreducens* colonized mice (**Figure 7C**). Within these samples, decreased prednisolone levels correlated with elevated 5β-dihydrocortisol and tetrahydrocortisol products concentrations, demonstrating steroid metabolism consistent with OsrABC pathway activity (**Figure 7C**). Notably, 3α,5β-tetrahydrocortisol, rather than 3β,5β-tetrahydrocortisol, was the predominant product detected in intestinal samples, contrasting with *C. steroidoreducens* cultures. This stereochemical difference may reflect enhanced *in vivo* expression of a *C. steroidoreducens* 3α-hydroxysteroid dehydrogenase or activity of a murine 3α-hydroxysteroid dehydrogenase previously reported in the intestinal epithelium.^52^

Analysis of systemic tissues revealed tissue-specific effects of *C. steroidoreducens* colonization on glucocorticoid metabolism. While hepatic and gallbladder prednisolone concentrations showed minimal change, serum levels were reduced 1.7-fold in *C. steroidoreducens-*colonized mice, accompanied by increases in 5β-dihydrocortisol and 3α,5β-tetrahydrocortisol (**Figure 7D**). These findings establish that *C. steroidoreducens* colonization can impact both local and circulating levels of glucocorticoid therapies, with potential implications for therapeutic efficacy in patients harboring elevated levels of steroid-metabolizing gut bacteria.

## Discussion

In this study, we characterize *Clostridium steroidoreducens* HCS.1, a previously uncharacterized gut bacterium that encodes a novel reductive pathway, OsrABC, for steroid hormone metabolism. Our work expands on previous studies of microbial steroid metabolism by demonstrating the widespread prevalence and activity of OsrB and OsrC, which catalyze the reduction of natural steroid hormones into their 3β,5β-tetrahydro derivatives. Notably, OsrB homologs are prevalent in Crohn’s disease-associated bacterial communities, implicating these enzymes in both health and disease contexts. Our gnotobiotic studies provide direct experimental validation that microbial steroid metabolism can substantially impact glucocorticoid bioavailability, establishing a mechanistic foundation for understanding therapeutic implications in inflammatory diseases.

A key aspect of this work is the link between microbial steroid metabolism and chronic inflammatory conditions, particularly Crohn’s disease. Our data indicate that *osrB* homologs are enriched in pro-inflammatory taxa such as *Clostridim_AQ innocuum* and *Ruminococcus_B gnavus*, which have previously been associated with Crohn’s disease.^49,50^ Pro-inflammatory gut microbes often exhibit a competitive advantage in inflammatory conditions and induce inflammation to generate conditions favorable for their growth.^51^ This suggests that these microbes may exploit steroid hormone metabolism to perpetuate inflammation, through the depletion of anti-inflammatory glucocorticoids.

The OsrABC pathway may have significant implications for glucocorticoid therapy, which remains a primary treatment for IBD.^46,47^ Our gnotobiotic studies demonstrate that *C. steroidoreducens* colonization markedly reduces prednisolone bioavailability, with 2-10-fold decreases in intestinal concentrations and approximately 2-fold reductions in serum levels. These findings establish that gut bacteria can efficiently convert therapeutic glucocorticoids into inactive 5β-reduced derivatives, directly compromising drug efficacy. The magnitude of these effects indicates that microbial metabolism represents a previously underappreciated mechanism contributing to steroid resistance in IBD patients. As such, patients harboring elevated levels of steroid-metabolizing bacteria may require personalized dosing regimens or alternative therapeutic approaches to achieve adequate immunosuppression and clinical response.

In sum, our study provides evidence that microbial steroid metabolism could have a clinically significant role shaping intestinal inflammation. The ability of gut bacteria to inactivate glucocorticoids may not only impact endogenous hormone signaling but also compromise the efficacy of steroid-based treatments in IBD. Future research should determine the extent to which microbial transformations contribute to steroid resistance and explore therapeutic strategies to prevent glucocorticoid degradation in IBD patients. Beyond the clinical considerations, our study highlights the broader significance of gut microbial steroid metabolism in human health. Notably, a recent publication independently identified the 3-oxo-Δ^4^-steroid hormone reductase activity of OsrB homologs, along with the characterization of additional novel gut bacterial enzymes that metabolize progestins.^53^ Together, these findings represent a step forward in delineating the broader landscape of microbial steroid hormone metabolism and its implications for human health.

## Resource Availability

### Lead Contact

Further information and requests for reagents may be directed to, and will be fulfilled by, the corresponding author, Sam Light

### Materials Availability

All chemicals used in this study were of analytical grade and were obtained as detailed in the Key Resources Table. Bacterial strains were sourced from the Duchossois Family Institute (DFI) Symbiotic Bacterial Strain Bank Repository (Chicago, IL) or other reputable repositories, as listed in the Key Resources Table.

### Data and Code Availability

The datasets generated in this study are available within the paper and its Supplementary Information. Transcriptomic datasets generated here can be accessed under BioProject PRJNA1139946. The whole genome sequences are accessible under accession SAMN42797975. Metagenomic data used in this study are publicly available from the Sequence Read Archive (SRA) or as supplementary data within this manuscript, under BioProject IDs PRJEB1220, PRJEB9576, PRJNA299502, PRJNA305507, PRJNA319574, PRJNA328899, PRJNA397112, PRJNA398089, PRJNA485056, PRJNA504891, PRJNA544527, and SRP057027.

NCBI accession codes for the studied proteins are MFU7515415.1 for OsrA, MFU7517346.1 for OsrB, and MFU7516964.1 for OsrC.

## Acknowledgements

Research reported in this publication was supported by funding from the National Institutes of Health (NIGMS R35GM146969 and NIDDK P30DK042086, via the University of Chicago Center for Interdisciplinary Study of Inflammatory Intestinal Disorders) and the Searle Scholars Program (to S.H.L), as well as the Deutsche Forschungsgemeinschaft (DFG, German Research Foundation) – Projektnummer 542537779 (to C.J.). We thank Dr. Eric Pamer and the Duchossois Family Institute Microbiome Metagenomics and Host-Microbe Metabolomics Facilities for advising and providing experimental support.

## Author Contributions

C.J., K.S., and S.H.L. conceived and designed the research. C.J., K.S., L.E., P.N.B., A.S.L., M.M., R.R., A.S., A.M.S., H.L., K.D.-T., B.H., and X.J. performed the experiments and analyzed the data. C.J., K.S., and S.H.L. wrote the manuscript with input from all authors.

## Experimental Model and Subject Details

### Animal Experiment

All mouse experiments were conducted in accordance with institutional guidelines and were approved by the University of Chicago’s Institutional Animal Care and Use Committee (IACUC) under protocol number 72617. Sixteen female germ-free

C57BL/6 mice (6-12 weeks old) were housed in sterile, autoclaved cages within a gnotobiotic facility under a 12-hour light/dark cycle with a controlled temperature (68-79 °F) and humidity (30-70%). The mice were provided with irradiated chow and acidified, autoclaved water ad libitum. Mono-colonization with *Clostridium steroidoreducens* HCS.1 was performed under sterile conditions two weeks prior to sample collection and colonization was confirmed after 7 days by fecal culturing and PCR detection of *osrB*. Following colonization, the mice were co-housed and handled using a sterile technique in a biosafety cabinet. Prednisolone was suspended in PBS and administered via oral gavage at a dose of 2 mg/kg body weight in a volume of 10 µL per gram of mouse weight. Fresh fecal pellets were collected immediately prior to sacrifice. The mice were euthanized, and samples were collected from the small intestine, cecum, colon, liver, gallbladder, and serum for downstream steroid analysis. Experiments shown in Figure 7 were performed once with the indicated number of biological replicates per group. Statistical analyses were conducted using two-sided Mann–Whitney U tests, followed by false discovery rate (FDR) correction using the Benjamini–Hochberg method. An FDR-adjusted *q* < 0.05 was considered significant.

## Method Details

### Steroid Enrichment Cultures

For each enrichment sample, 15 mM steroid hormone (cortisol, corticosterone, progesterone, or testosterone) suspensions were prepared in 1 mL basal growth medium, containing 19.2 g/L Na_2_HPO_4_·7H_2_O, 4.5 g/L KH_2_PO_4_, 0.75 g/L NaCl, 1.5 g/L NH_4_Cl, 2 g/L sodium acetate, 1.5 g/L sodium formate, 0.1 g/L tryptone, 0.1 g/L Bacto yeast extract, 1 g/L MgSO_4_, and a 1x stock of minerals and vitamins and was adjusted to pH 6.5. The 1x minerals and vitamins stock solution contained 0.1 mg/L FeCl_2_·4H_2_O, 0.846 mg/L MnSO_4_·H_2_O, 0.028 mg/L ZnSO_4_·7H_2_O, 0.148 mg/L CaCl_2_·2H_2_O, 0.002 mg/L CuSO_4_·5H_2_O, 0.002 mg/L CoCl_2_·7H_2_O, 0.002 mg/L H_3_BO_3_, 0.002 mg/L Na_2_MoO_4_·2H_2_O, 50 mg/L NaCl, 1.2 mg/L tri-sodium citrate, 0.05 mg/L biotin, 0.1 mg/L D-pantothenic acid, 0.05 mg/L lipoic acid, 0.1 mg/L niacinamide, 0.1 mg/L para-aminobenzoic acid, 0.1 mg/L pyridoxal HCl, 0.05 mg/L riboflavin, 0.1 mg/L thiamine HCl, and 0.01 mg/L vitamin B12. Homogenized fecal samples were pelleted and washed 3x in phosphate buffer saline (PBS), then resuspended in 1 mL PBS. 20 µL cell suspension was added to each enrichment culture condition and incubated for 72 hours. After 72 hours, 20 µL of each culture was used to inoculate fresh media supplemented with its respective compound. Cultures were passaged a total of 4 times. After the final passage, a portion of each condition was preserved in 20% glycerol and frozen at -80°C. The remaining culture was pelleted and processed for 16S rRNA sequencing.

Isolation of *Clostridium steroidoreducens* HCS.1 Preserved stocks of enrichment culture samples were plated onto fresh brain heart infusion (BHI) agar and incubated for 4 days at 37°C under anaerobic conditions (5% H_2_, 10% CO_2_, 85% N_2_). Distinct colonies were passaged to confirm purity and identified by 16S rRNA V4-V5 variable region sequencing. Purified isolates of HCS.1 were stored at -80°C in a 20% glycerol suspension. Frozen glycerol stocks were deposited in the DFI Symbiotic Bacterial Strain Bank Repository (https://dfi.cri.uchicago.edu/biobank/).

### Steroid Clearance Assay

To prepare steroid clearance assays, progesterone and cortisol amounts for a final concentration of 12 mM or 16 mM, respectively, were sterilized by suspension in 70% v/v ethanol, followed evaporation at room temperature for 2 hours. Dried steroid powders were sifted into autoclaved BHI agar and stirred rapidly, shortly before pouring into plates. Solid plates were stored under anaerobic conditions at 25°C for at least 24 hours prior to use.

To test HCS.1 steroid clearance, solid BHI plates were incubated at 37°C for 2 days. After 2 days, single colonies were picked and suspended in 200 μL PBS. Aliquots of the cell suspension were spread onto solid steroid plates and incubated anaerobically for 5 days at 37°C. To test steroid clearance of other bacterial strains (detailed in **Dataset S3**), solid BHI plates were incubated at 37°C for 4 days. After 4 days, single colonies from each isolate were picked and suspended in 200 μL PBS. 2 μL aliquots were spotted onto solid progesterone plates, dried, and incubated anaerobically for 3 days at 37°C.

### 16S rRNA Gene Sequencing

Cells from final steroid enrichment passages were collected by centrifugation and their genomic DNA extracted using the QIAamp PowerFecal Pro DNA kit (Qiagen). Briefly, samples were suspended in a bead tube (Qiagen) along with lysis buffer and loaded on a bead mill homogenizer (Fisherbrand). Samples were then centrifuged, and the supernatant was resuspended in a reagent that effectively removed inhibitors. DNA was then purified routinely using a spin column filter membrane and quantified using Qubit. The 16S rRNA variable V4-V5 region was amplified using universal bacterial primers, 564F and 926R. Amplicons were purified using magnetic beads, then quantified and pooled at equimolar concentrations. The Qiagen QIAseq one-step amplicon library kit was used to ligate Illumina sequencing-compatible adaptors onto amplicons. Reads were sequenced on an Illumina MiSeq platform to generate 2 x 250 base pair reads, with 5,000-10,000 reads per sample. Amplified 16S rRNA amplicons were processed using the DADA2 pipeline in R. Forward reads were trimmed at 210 bp and reverse reads at 150 bp, to remove low-quality nucleotides. Chimeras were detected and removed using default parameters. Amplicon sequence variants between 300 and 360 bp in length were taxonomically assigned to the genus level using the RDP Classifier (v2.13) with a minimum bootstrap confidence score of 80.

### Whole Genome Sequencing

To prepare HCS.1 for whole genome sequencing, 10 mL BHI broth was inoculated with cells from a single bacterial colony and incubated anaerobically at 37°C for 48 hours. The culture was centrifuged at 4000 x g for 10 minutes. The resulting pellet was resuspended, washed in phosphate buffer saline (PBS), and re-centrifuged. Samples for Illumina short sequencing were extracted using the QIAamp PowerFecal Pro DNA kit (Qiagen), as described in the preceding subsection. Libraries were prepared using 200 ng of genomic DNA using the QIAseq FX DNA library kit (Qiagen). Briefly, DNA was fragmented enzymatically into shorter fragments and desired insert size was achieved by adjusting fragmentation conditions. Fragmented DNA was end repaired and ‘A’s’ were added to the 3’ends to stage inserts for ligation. During ligation step, Illumina compatible Unique Dual Index (UDI) adapters were added to the inserts and prepared library was PCR amplified. Amplified libraries were cleaned up, and QC was performed using Tapestation 4200 (Agilent Technologies). Libraries were sequenced on an Illumina NextSeq 1000/2000 to generate 2×150bp reads.

Samples for Nanopore and Illumina hybrid assemblies were extracted using the high molecular weight NEB Monarch Genomic DNA Purification Kit. DNA quality was assessed with the Agilent Tapestation 4200. Nanopore libraries were prepared using the Rapid Sequencing Kit (SQK-RAD114) and sequenced on MinION R10.4.1 flow cells. Nanopore reads were base-called using ONT Guppy basecalling software version 6.5.7+ca6d6af, minimap2 version 2.24-r1122, and was demultiplexed using ONT Guppy barcoding software version 6.5.7+ca6d6af using local HPC GPU. N50 of the nanopore long read is 7077 base pairs, the average read length is 4529.4 base pairs, while the average read quality is 15.6, which is typical of Nanopore reads. Hybrid assembly was performed with both nanopore and Illumina short reads using Unicylcer v0.5.0.^54,55^

### Genome Classification

The classification of strain HCS.1 as a novel species, *Clostridium steroidoreducens* sp. nov. was performed using GTDB-Tk (version 2.3.2)^56^ on the KBase platform. Genome quality assessment, phylogenetic placement, and taxonomic classification were performed according to GTDB guidelines. The HCS.1 genome was uploaded to the KBase website and analyzed using the GTDB-Tk classify workflow, which assigns genomes to the closest known taxa based on conserved marker genes. Strain HCS.1 was classified within the genus *Clostridium* but did not match any known species in the Genome Taxonomy Database (GTDB, version r207). Phylogenetic placement within the GTDB bacterial tree confirmed that the strain represented a distinct lineage, supporting its designation as a new species.

### Transcriptomics of *Clostridium steroidoreducens* HCS.1

To prepare HCS.1 samples for transcriptomic analysis, six 50 mL Lysogeny broth cultures were inoculated with bacterial cells and shaken anaerobically at 37°C for 48 hours. After 48 hours, cortisol powder was added to 3 cultures, to a final concentration of 8 mM. All 6 cultures were incubated for an additional 4 hours, then pelleted at 4000 x g for 10 minutes. The resulting pellets were flash-frozen in a dry ice/ethanol bath and stored at -80°C until ready for subsequent processing. Cell pellets were thawed and total RNA from biological replicates were extracted using the Maxwell RSC instrument (Promega). Extracted RNA was quantified using Qubit, and integrity was measured using TapeStation (Agilent Technologies). Libraries from ribo-depleted samples were constructed using the NEB’s Ultra Directional RNA library prep kit for Illumina. First, up to 500 ng total RNA was subjected to ribosomal RNA depletion (for bacteria) using NEBNext rRNA depletion kit. Ribosomal -RNA depleted samples were fragmented based on RNA integrity number (RIN). Post cDNA synthesis, Illumina compatible adapters were ligated onto the inserts and the final libraries were quality assessed using TapeStation (Agilent technologies). Libraries were normalized using library size and final library concentration (as determined by Qubit). Library concentration (ng/ul) was converted to nM to calculate dsDNA library concentration. Equimolar libraries were then pooled together at identical volumes to ensure even read distribution across all samples. Normalized libraries were then sequenced on Illumina’s NextSeq 1000/2000 at 2x100bp read length.

High-quality reads were mapped to the circularized hybrid assembled genome of HCS.1 (NCBI: CP170704), using Bowtie2 (v.2.4.5), and sorted with Samtools (v1.6). Read counts were generated using featureCounts (v2.0.1) with Bakta annotations.^57^ Gene expression was quantified as the total number of reads uniquely aligning to the reference genome, binned by annotated gene coordinates. Differential gene expression and quality control analyses were performed using DESeq2 in R with Benjamini–Hochberg false discovery rate adjustment applied for multiple testing corrections.^58^

### Molecular Cloning and Plasmid Construction

Gene transformations were performed by Gibson assembly using 2x NEBuilder® HiFi DNA Assembly Master Mix (New England Biolabs, NEB, E2621X). Primers were designed using SnapGene (see **Table S1**), incorporating 20 bp flanking regions complementary to a linearized expression vector (pMCSG53) and the gene of interest from the HCS.1 genome. PCR, cloning, and transformation were performed according to the protocols provided on the NEB website. The Gibson assembly reaction was incubated at 50°C for 1 hour and then transformed into *E. coli* XL1-Blue competent cells according to the manufacturer’s protocol. Transformed cells were plated on Luria-Bertani (LB) agar plates containing 100 µg/mL carbenicillin, and successful transformations were confirmed using sequencing primers specific for the backbone vector (see **Table S1**). Positive colonies were validated and sequenced by the University of Chicago Genomics Facility. The final constructs were then transformed into chemically competent *E. coli* Rosetta™ (DE3) competent cells (Novagen) according to NEB protocols. Transformed cells were plated on LB agar plates supplemented with 100 µg/mL carbenicillin.

### Heterologous Protein Production in *E. coli*

Protein production in *E. coli* Rosetta cells was performed under aerobic conditions for all short-chain dehydrogenases (SDRs) and under anaerobic conditions for OsrA and OsrB. Cultures were grown in either 2x YT medium (20 g/L tryptone, 10 g/L yeast extract, and 5 g/L NaCl) or TB medium (12 g/L tryptone, 24 g/L yeast extract, 4 mL/L glycerol, 9. 4 g/L K_2_HPO_4_, and 2.2 g/L KH_2_PO_4_) supplemented with 0.5% (w/v) glucose and 1 mM ferric ammonium citrate, respectively. Induction of protein expression was initiated during the exponential phase when optical densities (OD_600_) reached 0.4-0.6 by the addition of 1 mM isopropyl β-D-1-thiogalactopyranoside (IPTG). Cultures were incubated for 3-5 hours at 37°C with shaking at 200 rpm. Cells were harvested by centrifugation at 4500 × g for 20 minutes. Cell pellets were frozen at -80°C for storage prior to subsequent experimental use. Protein production was confirmed by SDS-PAGE analysis.

### Protein Purification

Frozen cell pellets were supplemented with 0.1 mg/mL DNase and lysed either aerobically (for SDRs) or anaerobically (for OsrA and OsrB) using a Thermo Spectronic French pressure cell at 1,100 PSI. The crude cell extract was centrifuged at 75,600 x g for 30 minutes. The supernatant was filtered through a 0.2 µm nylon membrane (Fisher Scientific) before being applied to a purification system.

For SDRs the filtered extract was applied to an ÄKTA pure system (Cytiva) using a 1 mL Strep-Tactin®XT 4Flow® column (Iba Lifesciences). The column was equilibrated with 10 volumes of equilibration buffer (100 mM Tris/HCl, pH 8.0, 150 mM NaCl) at 1 mL/min and 8°C. Proteins were loaded via a 5 mL loop, followed by washing of non-specifically bound proteins. Elution was performed with 5 mL elution buffer (100 mM Tris/HCl, pH 8.0, 150 mM NaCl, and 50 mM biotin). Eluted proteins were collected in 1 mL fractions and were concentrated using Pierce™ Protein concentrators (10 kDa), desalted, and either used directly or transferred to storage buffer (50 mM Tris/HCl, pH 7.5, 10% (w/v) glycerol, and 50 mM NaCl) using PD-10 desalting columns (Cytiva) before storage at -80°C.

OsrA and OsrB purification was performed in an anaerobic chamber. A Strep-Tactin®XT 4Flow® gravity column (Iba Lifesciences) was used with a WET FRED system (Iba Lifesciences) to maintain a constant flow rate of ∼1 mL/min, adjusted using a lab jack stand (LABALPHA). The column was equilibrated with anaerobic equilibration buffer (100 mM Tris/HCl, pH 8.0, 150 mM NaCl) and elution was performed with anaerobic elution buffer (100 mM Tris/HCl, pH 8.0, 150 mM NaCl, and 50 mM biotin). Eluted proteins were collected in 1 mL tubes, concentrated and desalted using Pierce™ Protein Concentrators PES (10K MWCO, 0.5 mL) at 10,000 × g in a microcentrifuge. Proteins were either used directly for enzymatic assays or transferred to anaerobic storage buffer (50 mM Tris/HCl, pH 7.5, 10% (w/v) glycerol, and 50 mM NaCl) and frozen at -80°C.

### In Vitro Enzyme Assays

The activities of purified SDRs were determined using reaction mixtures in 96-well plates with a total volume of 100 µL. The reaction mixture contained 50 mM phosphate buffer (pH 7.0), 0.2 mM NADPH, 100 µM steroids (from 2 mM stock solutions in methanol), and 1 to 100 µg/mL protein, depending on the enzyme activity. Enzyme activity was monitored by measuring the reduction of NADPH at 340 nm using a Cytation 5 imaging reader (BioTek) at 25°C. A NADPH standard curve was analyzed under identical conditions with an extinction coefficient of 1586 M^-1^ for quantitation.

The substrate preferences of OsrA and OsrB were analyzed under anaerobic conditions using a 100 µL reaction mixture containing 50 mM phosphate buffer (pH 7.0), 1 mM methyl viologen, 0.2 mM sodium dithionite, 0.1 mM steroids (from 2 mM stock solutions in methanol), and 0.1 to 1 mg/mL protein. Enzyme activity was monitored by measuring electron donor reduction at 605 nm using an EPOCH 2 microplate reader (BioTek) at 37°C. Quantification was performed using an electron donor (methyl viologen) standard curve generated under the same conditions with an extinction coefficient of 2136 M^-1^ for quantitation.

### Whole-cell Assays in Bacterial Cultures

A complete list of strains used in this study is provided in the Key Resources Table. Strains were incubated under anaerobic conditions (85% N_2_, 10% CO_2_, 5% H_2_) at 37°C in an anaerobic chamber (Coy Laboratory). Liquid brain-heart infusion (BHI) broth supplemented with 100 µM steroids (prepared from 10 mM stock solutions in methanol) was used for growth. Cultures were inoculated at 1% (v/v) from a pre-culture and incubated anaerobically. Steroids were added after 4 hours of growth during the exponential phase. In the case of heterologously expressed enzymes in *E. coli*, 1 mM IPTG was added simultaneously to induce protein expression. Bacterial cultures were extracted by adding 50 volumes of methanol containing 0.5 µM methylprednisolone as an internal standard for LC-MS-QTOF analysis.

### LC-MS-QTOF Analysis

Extracted samples were vortexed and centrifuged twice at 21,000 × g for 15 minutes, with the supernatant transferred to new tubes after each centrifugation step. The methanol fraction was filtered through 0.2 µm nylon membrane filters prior to LC-MS analysis. Samples were analyzed using an Agilent 6540 UHD Q-TOF mass spectrometer coupled to an Agilent 1290 UHPLC system system (Agilent Technologies). Separation was performed on a XBridge C18 column (2.1×100mm, 3.5 µm particle size) using 0.1% aqueous formic acid and acetonitrile with 0.1% formic acid as mobile phases. The separation gradient ranged from 20% to 100% acetonitrile over 4 minutes at 50°C with a flow rate of 0.5 mL/min. Mass spectra were acquired in negative ion mode for glucocorticoids ([M+FA-H]^-^) or positive ion mode for all other steroids ([M+H]^+^), with an ion spray voltage of 3500 V and a nozzle voltage of 2000 V. The source temperature was set to 300°C, and the gas flow rate was 12 L/min. Data were processed and visualized using MassHunter software version 10.

### LC-MS-TQ Analysis of Tissues

For total glucocorticoid analysis, gall bladder, liver, fecal, cecal, and intestinal samples were extracted in 80:20 methanol/water at a volume of 10 µL per mg sample, containing 0.5 µM methylprednisolone as an internal standard. Serum (50 µL each) were extracted in nine volumes of the same solvent mixture. Samples were homogenized at 4°C using a Bead Mill 24 (program 1: S = 2.10, T = 1:00, C = 10, D = 0:01, Fisherbrand), followed by centrifugation at 21,000 × g for 5 minutes. The resulting supernatants were collected and evaporated to dryness using a Genevac EZ-2 Series evaporator (ATS Scientific Products) operated in aqueous mode. Residues were reconstituted in 100 µL of 2000 U/mL β-glucuronidase (from *Helix pomatia*) in 0.1 M sodium acetate buffer (pH 5.0) and incubated at 37 °C for 2 hours. Steroids were subsequently extracted with 400 µL methyl tert-butyl ether (MTBE) and dried using the Genevac EZ-2 Series in very low boiling point mode. Samples were then reconstituted in 10 µL per mg sample (solid samples) or 10 µL per µL sample (serum) of methanol containing 0.2 µM d4-cortisol, centrifuged at 21,000 × g for 5 minutes, and the supernatants were filtered through a 0.2 µm nylon filter prior to LC–MS analysis.

Glucocorticoid quantification was performed on an Agilent 6460 Triple Quadrupole LC/MS system coupled to a 1290 UHPLC (Agilent Technologies) operated in negative electrospray ionization mode. Steroids were separated on a XBridge C18 column (2.1×100mm, 3.5 µm particle size) using a linear gradient (0.4 mL/min) of water with 0.02% formic acid and acetonitrile with 0.02% formic acid, ramping from 5% to 60% acetonitrile over 4 minutes at 40°C. The following transitions in dynamic MRM mode were monitored: prednisolone (m/z 405.2 → 329.2), cortisol (407.2 → 331.2), dihydrocortisol (409.2 → 333.2), and tetrahydrocortisol (411.2 → 355.2), all detected as [M+FA-H]^−^ adducts. Instrument settings were as follows: ion spray voltage 3500 V, nozzle voltage 500 V, gas temperature 300°C, and gas flow 12 L/min.

### Phylogenetic Tree Construction

Genome metadata were retrieved from a local UHGG database and used to map genome IDs to species names. A comparative analysis was performed using BLASTp (version 2.15.0+) to search for homologs of the target sequence against the UHGP-100 database, limiting results to the top 5000 hits. The BLASTp output was processed to map genome IDs to species names and format the sequences in FASTA format, removing duplicates to ensure data quality. Additional sequences were appended to the data set as needed. Sequence alignment was performed using Clustal Omega (version 1.2.2)^59^ with output formatted as FASTA. Header sanitization was performed to remove special characters, and duplicate sequences were filtered out using Python (v3.12). Phylogenetic analysis was performed using IQ-TREE (version 2.3.6)^60^ with automatic model selection to determine the best-fitting substitution model based on the data. The reliability of the phylogenetic trees was assessed using 1,000 ultrafast bootstrap replicates to assess branch support. The final phylogenetic trees were visualized and interpreted using the Interactive Tree of Life (iTOL)^61^ to explore the evolutionary relationships among the identified protein sequences.

### Metagenomic Analysis

To identify relevant sequences of OsrA or OsrB, a BLASTp (version 2.15.0+) search was first conducted against the UHGP-100 database, applying a 49% similarity threshold based on experimental evidence. For OsrB, the identified protein sequences were used to construct a phylogenetic tree using Clustal Omega for alignment and IQ-TREE with the “mtest” model selection and 1,000 bootstrap replicates as described above. Based on the initial analysis, 25 sequences that were not phylogenetically related to OsrB were manually excluded so that only sequences relevant to the target enzymes were retained for downstream analysis.

After phylogenetic filtering, genome information was traced back using the UHGP protein IDs. The corresponding genomes were downloaded from the Unified Human Gastrointestinal Genome (UHGG) database using FTP links provided in the metadata file. The genomes were then used for further analysis, where each protein sequence was screened against the respective genome using tblastn with a e-value threshold of 1e^-200^. The best nucleotide sequence was selected for each protein based on coverage and bit score and subsequently compiled into a combined FASTA file.

Metagenomic samples were downloaded from the Sequence Read Archive (SRA). Reads were quality trimmed to remove adapter sequences using TrimGalore with default settings,^62^ and potential human contamination was removed by mapping the reads to the human reference genome (T2T-CHM13v2.0) using Bowtie2 (version 2.5.3) and removing the mapped reads with Samtools (version 1.61.1).^63,64^ Samples were then mapped to the gene reference datasets for *osrA* and *osrB* using Bowtie2 (version 2.5.3), and copies per million (CPM values) were calculated for each gene in each sample. Samples with total read counts below 1,000,000 were excluded.

Normality was assessed using the Shapiro-Wilk test; if both groups were normal (p > 0.05), a two-sided Welch’s t-test was used. If normality was not met, a two-sided Mann-Whitney U test was applied. Statistical analyses were conducted in Python (v3.12) with libraries including Pandas (v2.2.0), NumPy (v1.26.4), Matplotlib (v3.8.3), Seaborn (v0.13.2), and Pingouin (v0.5.5).

