## Supplementary material for "Gut Bacteria Metabolize Natural and Synthetic Steroid Hormones via the Reductive OsrABC Pathway": Figure S

**Figures S1-S6, Tables S1**

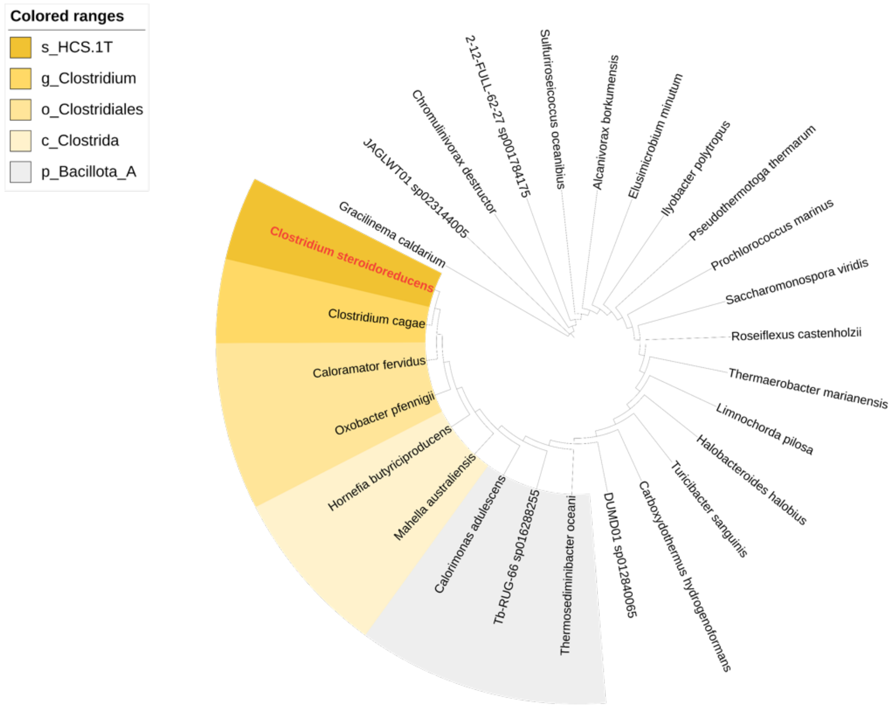


**Figure S1. Phylogenetic analysis supporting the assignment of HCS.1 to the *Clostridium* genus.** The tree was generated from an alignment of conserved marker genes.

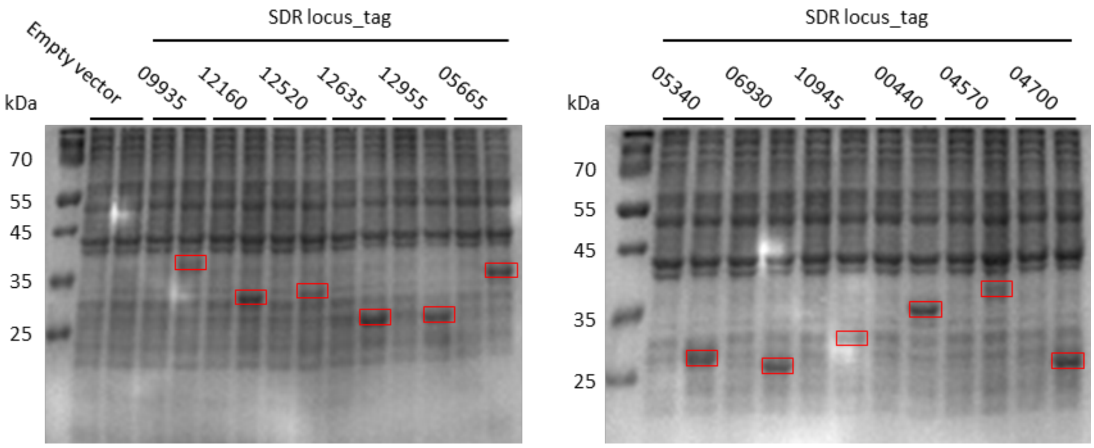


**Figure S2. 12% SDS-PAGE of *E. coli* lysates expressing SDR domain-containing proteins.** SDS-PAGE analysis showing the heterologous production of *C. steroidoreducens* SDR domain-containing proteins in *E. coli*, pre- and post-isopropyl β-D-1-thiogalactopyranoside (IPTG) induction. Red boxes highlight expressed protein.

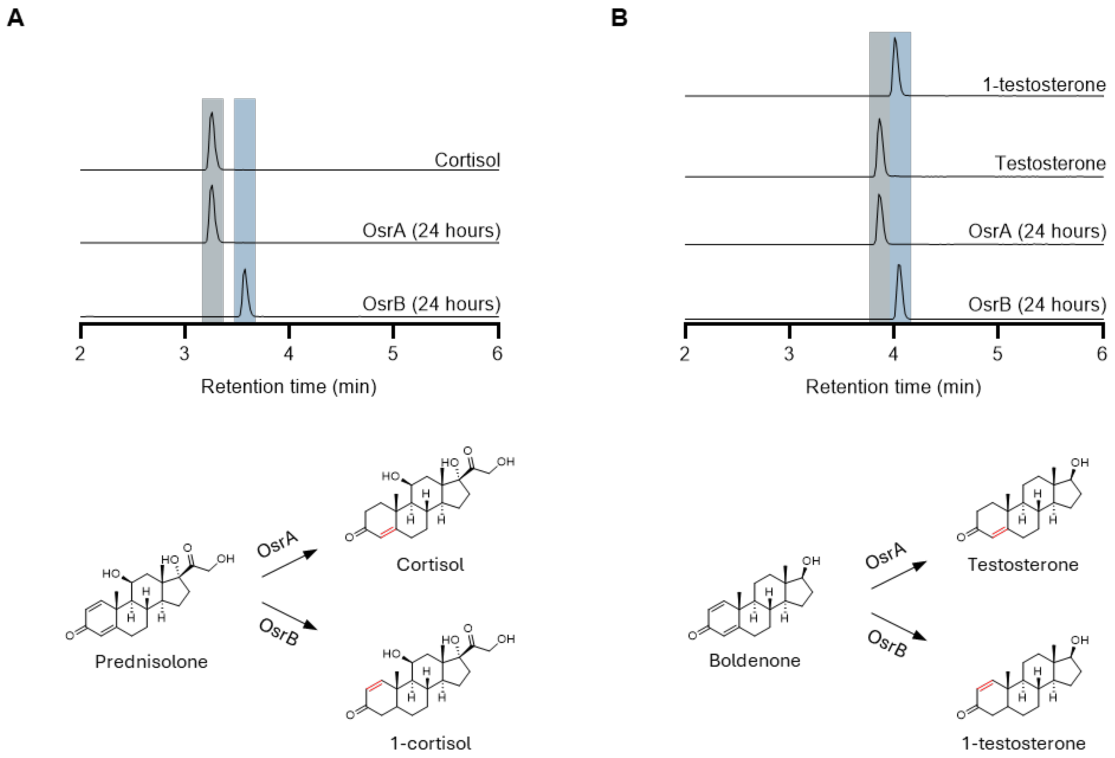


**Figure S3. OsrA and OsrB products from Δ$^1$- and Δ$^4$-steroid hormone substrates.** (A) Products following prednisolone incubation with purified OsrA or OsrB. Comparison to a cortisol reference standard confirm that OsrA generates the Δ$^1$-reduced product and show that OsrB produces a distinct cortisol isomer. (B) Products following boldenone incubation with purified OsrA or OsrB. Comparison to reference standards confirm that OsrA and OsrB generate Δ$^1$- and Δ$^4$-reduced products, respectively.

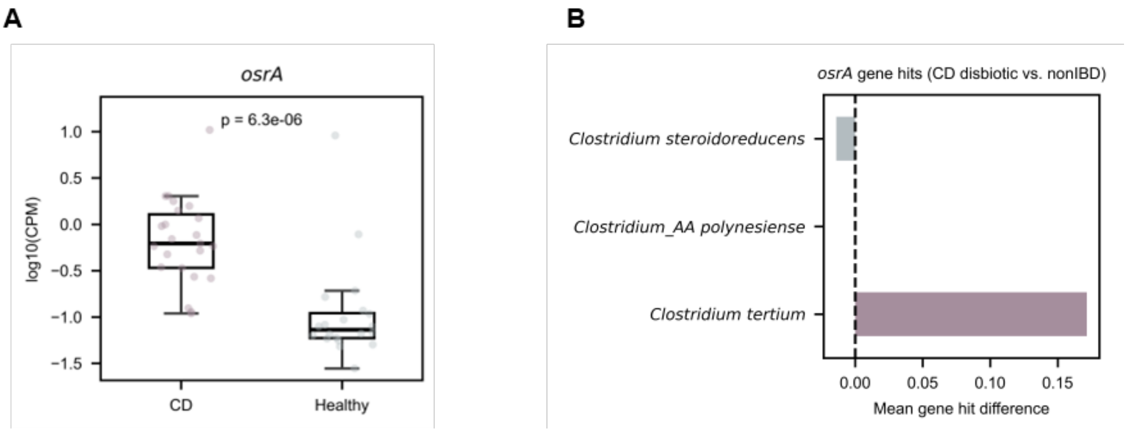


**Figure S4. *osrA* levels in expanded dataset of healthy and Crohn's disease metagenomes.** (A) Reads mapping to *osrA* homologs in expanded dataset of Crohn's disease (CD) patient metagenomes relative to healthy controls. (B) Difference in *osrA* homolog levels in CD relative to healthy metagenomes. Sample sizes: for $n_{\text{Healthy}}$ = 20 and $n_{\text{CD}}$ = 15.

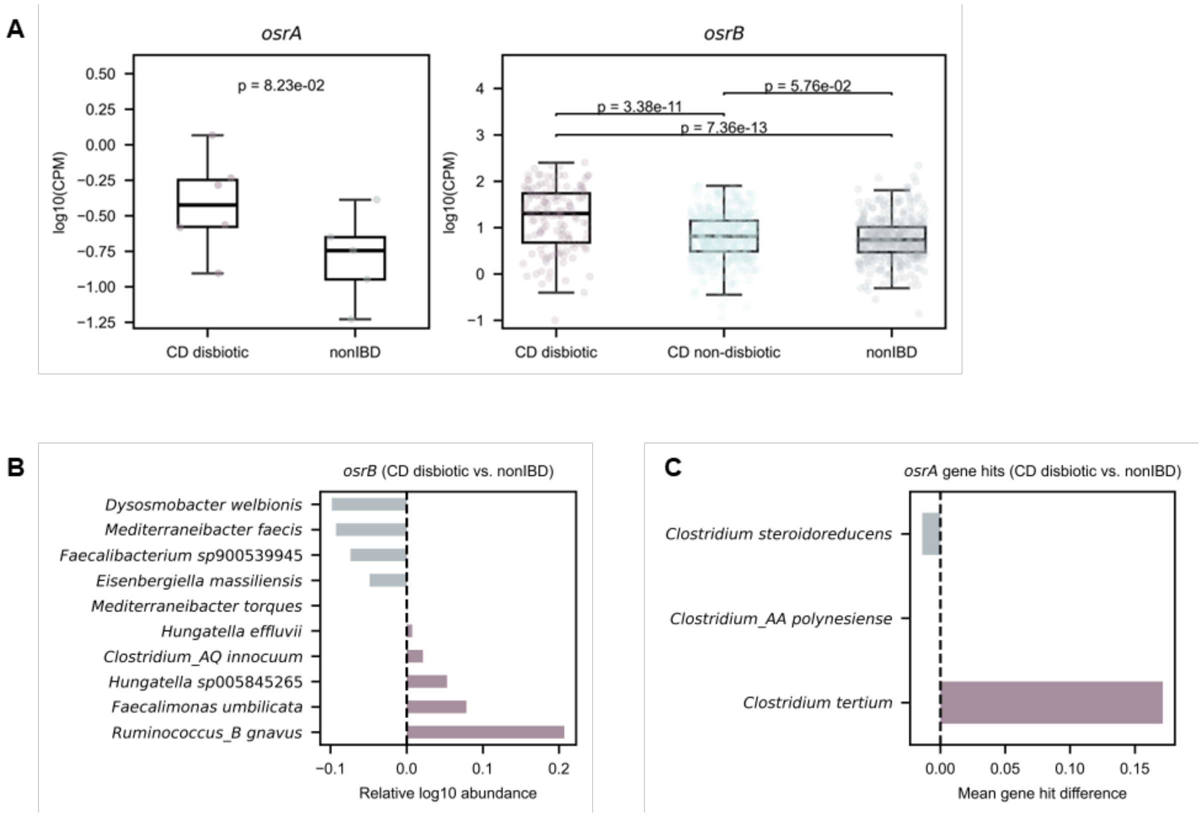


**Figure S5. Association of *osrA* and *osrB* with Crohn's disease in Integrative Human Microbiome Project metagenomes.** (A) Read counts mapping to *osrA* and *osrB* homologs in Crohn's disease (CD) metagenomes relative to non-IBD controls. CPM stands for copies per million. Sample sizes: for osrA, $n_{\text{CD disbiotic}}$ = 6 and $n_{\text{nonIBD}}$ = 5; for osrB, $n_{\text{CD disbiotic}}$ = 143, $n_{\text{non-disbiotic}}$ = 433 and $n_{\text{nonIBD}}$ = 355. (B) Difference in *osrB* homolog levels among taxa with the most significant relative abundance changes between CD dysbiotic and non-IBD metagenomes. (C) Difference in *osrA* homolog levels in CD dysbiotic relative to non-IBD metagenomes.

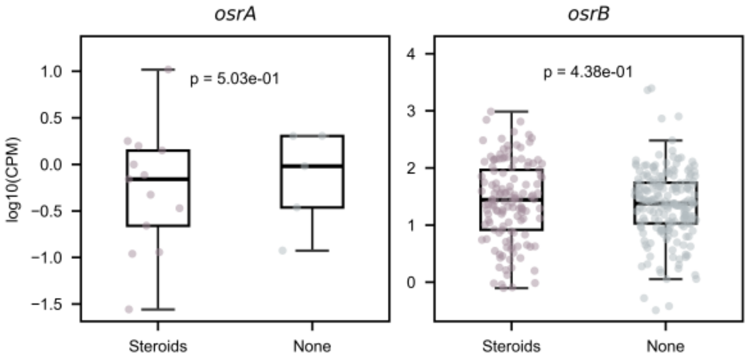


**Figure S6. Association of *osrA* and *osrB* with steroid usage in the Lewis et al study.** Read counts mapping to *osrA* and *osrB* homologs in Crohn's disease (CD) patient metagenomes, grouped by glucocorticoid treatment status. Sample sizes: for osrA, $n_{\text{Steroids}}$ = 13 and $n_{\text{None}}$ = 5; for osrB, $nS_{teroids}$ = 122 and $n_{\text{None}}$ = 176.

**Table S1: Primers used in this study.**

| Gene/Identifier | Primer | Sequence (5' --> 3') | Length |
|---|---|---|---|
| *osrA* (nStrep) | for | TTTAAGAAGGAGATATACATATGTGGAGCCACCCGCAGTTCGAAAAGGGAGGAGGAGGAGGAGGAAAATTATTTAATGATGGACATATTGGATCATTATC | 100 |
| | rev | GCTTTGTTAGCAGCCGGATCTTATATTTCTATTCCTACTTGGAAACCTTCAG | 52 |
| *osrB* (nStrep) | for | TTTAAGAAGGAGATATACATATGTGGAGCCACCCGCAGTTCGAAAAGGGAGGAGGAGGAGGAGGAGGAGGAGCGTTTGAGAATGTTTTTAGTCC | 94 |
| | rev | GCTTTGTTAGCAGCCGGATCCTAAATTTGTCTTGCTACATTTACTGC | 47 |
| *osrC* *ACFYH6_05340* (nStrep) | for | TTTAAGAAGGAGATATACATATGTGGAGCCACCCGCAGTTCGAAAAGGGAGGAGGAGGAAGCGGAGGAGGAGGAAGCGCTCGTTTAAAAGGGAAAGTAGC | 100 |
| | rev | GCTTTGTTAGCAGCCGGATCTTATAAAGTACTACCAAAATCTGAAATTATAGTTTG | 56 |
| *ACFYH6_12520* (nStrep) | for | TTTAAGAAGGAGATATACATATGTGGAGCCACCCGCAGTTCGAAAAGGGAAGCGGAAGCGGAAGCAAACTTTGTGGTAAGGTAGCC | 86 |
| | rev | GCTTTGTTAGCAGCCGGATCTTAAAATGCCGTCCAACCTG | 40 |
| *ACFYH6_09935* | for | TTTAAGAAGGAGATATACATATGAATACAGAGTTTACAAATAAAATTCCAG | 51 |
| | rev | GCTTTGTTAGCAGCCGGATCTTATGTTGTTACATTTGCTCCTCC | 44 |
| *ACFYH6_12160* | for | TTTAAGAAGGAGATATACATATGGAACGTTTAAAAGGTAAAGTTG | 45 |
| | rev | GCTTTGTTAGCAGCCGGATCTTACATTGCAGTTAATCCACCATC | 44 |
| *ACFYH6_12520* | for | TTTAAGAAGGAGATATACATATGAAACTTTGTGGTAAGGTAGC | 43 |
| | rev | GCTTTGTTAGCAGCCGGATCTTAAAATGCCGTCCAACCTG | 40 |
| *ACFYH6_12635* | for | TTTAAGAAGGAGATATACATATGAGAGTAGCATTAATAACTGGAG | 45 |
| | rev | GCTTTGTTAGCAGCCGGATCTTACTCTTCATAAATCATCTTTTTTGTCATTCC | 53 |
| *ACFYH6_12955* | for | TTTAAGAAGGAGATATACATATGGTTGTTAATAAAAATGCAATAGTCAC | 49 |
| | rev | GCTTTGTTAGCAGCCGGATCTTAATGCATAACCATTCCACCATC | 44 |
| *ACFYH6_05665* | for | TTTAAGAAGGAGATATACATATGAAAAATGTAGTAATAACAGGAAGTACTC | |
| | rev | GCTTTGTTAGCAGCCGGATCTTATTTTAAAAGTTCTCTCTTCTTAAAAGAAGAC | |
| *osrC* *ACFYH6_05340* | for | TTTAAGAAGGAGATATACATATGGCTCGTTTAAAAGGGAAAG | 42 |
| | rev | GCTTTGTTAGCAGCCGGATCTTATAAAGTACTACCAAAATCTGAAATTATAGTTTG | 56 |
| *ACFYH6_06930* | for | TTTAAGAAGGAGATATACATATGGAACGTTTAAAAGATAAAGTTGC | 46 |
| | rev | GCTTTGTTAGCAGCCGGATCTTACATTGCAGTTAATCCACCATC | 44 |
| *ACFYH6_10945* | for | TTTAAGAAGGAGATATACATATGGAACGTTTAGAAGGAAAAATAG | 45 |
| | rev | GCTTTGTTAGCAGCCGGATCCTAAATCATTGCTCCATCAATTTTAATTATTTG | 53 |

| | | | |
|---|---|---|---|
| *ACFYH6_00440* | for | TTTAAGAAGGAGATATACATATGGGTAAGCCGTATGGTAG | 40 |
| | rev | GCTTTGTTAGCAGCCGGATCCTAACCTTTGTTCATCATTAATATTATTTT TCTAG | 55 |
| *ACFYH6_04570* | for | TTTAAGAAGGAGATATACATATGTTCGTTAGAATTACAAATGGTTTG | 47 |
| | rev | GCTTTGTTAGCAGCCGGATCTTACCAAGTACGAGATTTCATTTCTTTG | 48 |
| *ACFYH6_04700* | for | TTTAAGAAGGAGATATACATATGTTAAGATTAGAAAATAAGGTTGCTA TAG | 51 |
| | rev | GCTTTGTTAGCAGCCGGATCCTATAATCTCATTCCACCATTAAC | 44 |
| Linearized plasmid | pMCS_for | ATGTATATCTCCTTCTTAAAGTTAAACAAAATTATTTC | 38 |
| | pMCS_rev | GATCCGGCTGCTAACAAAGC | 20 |
| Sequencing Primers | Seq_for | TAATACGACTCACTATAGGGGAATTG | 26 |
| | Seq_rev | CAAAAAACCCCTCAAGACCC | 20 |
